# Soldier neural architecture is temporarily modality-specialized but poorly predicted by repertoire size in the stingless bee *Tetragonisca angustula*

**DOI:** 10.1101/2021.08.15.456373

**Authors:** Kaitlin M. Baudier, Meghan M. Bennett, Meghan Barrett, Frank J. Cossio, Robert D. Wu, Sean O’Donnell, Theodore P. Pavlic, Jennifer H. Fewell

**Affiliations:** School of Biological, Environmental and Earth Sciences, The University of Southern Mississippi, Hattiesburg, MS, USA; School of Life Sciences, Social Insect Research Group, Arizona State University, Tempe, AZ, USA; USDA-ARS Carl Hayden Bee Research Center, Tucson, AZ, USA; Department of Biology, Drexel University, Philadelphia, PA, USA; Department of Biodiversity, Earth and Environmental Science, Drexel University, Philadelphia, PA, USA; School of Computing, Informatics, and Decision Systems Engineering, Arizona State University, Tempe, AZ, USA; School of Sustainability, Arizona State University, Tempe, AZ, USA; School of Complex Adaptive Systems, Arizona State University, Tempe, AZ, USA

**Keywords:** Abejas angelitas, caste, division of labor, group defense, social insects, temporal polyethism

## Abstract

Individual heterogeneity within societies provides opportunities to test hypotheses about adaptive neural investment in the context of group cooperation. Here we explore neural investment in defense specialist soldiers of the eusocial stingless bee (*Tetragonisca angustula*) which are age sub-specialized on distinct defense tasks, and have an overall higher lifetime task repertoire than other sterile workers within the colony. Consistent with predicted behavioral demands, soldiers had higher relative visual (optic lobe) investment than non-soldiers but only during the period when they were performing the most visually demanding defense task (hovering guarding). As soldiers aged into the less visually demanding task of standing guarding this difference disappeared. Neural investment was otherwise similar across all colony members. Despite having larger task repertoires, soldiers had similar absolute brain size and smaller relative brain size compared to other workers, meaning that lifetime task repertoire size was a poor predictor of brain size. Together, our results are consistent with the specialized but flexible defense strategies of this species, broadening our understanding of how neurobiology mediates age and morphological task specialization in highly cooperative societies.

## Introduction

Cognitive investment within highly cooperative social groups can be distributed across individuals [1], and the question of how differences in brain investment among heterogeneous individuals within social groups relates to adaptive group function is a popular area of study [2–4]. Eusocial insect colonies often have pronounced task and/or morphological differentiation among colony members [5], making them excellent models for studying the effects of behavioral heterogeneity on the interplay between individual neural investment and socially coordinated function.

Among eusocial insects, the most commonly evolved category of morphologically specialized workers are soldiers: large-bodied non-reproductives performing colony defense behaviors [6]. The most well-studied soldiers are perhaps those of termites and ants [6–10], but morphologically distinct soldiers have also evolved multiple times among eusocial stingless bees [11]. Unlike ant soldiers, morphologically distinct soldiers of the stingless bee, *Tetragonisca angustula* [12] perform all of the tasks that non-soldier workers (henceforth “minors”) perform but on an accelerated age-trajectory, switching to a repertoire dominated by colony defense tasks in the last two weeks of life (Figure 1) [13]. Soldiers of *T. angustula* further sub-specialize among different end-of-life defense tasks according to age [14]. Younger hovering guards primarily protect against heterospecific invasion using visual and volatile chemical cues while older standing guards on the nest entrance tube also intercept conspecific non-nestmates using close-range olfaction of non-volatile chemical cues [14–18]. These tasks place different demands on visual and olfactory acuity and processing among these different soldier age groups. There is behavioral evidence for specialization in olfactory invader cue detection between soldier types (hovering versus standing guards) [19] which is not associated with differential antennal sensitivity [20]. Neural tissue is energetically costly to produce and maintain, and so neural tissue is expected to be lower in volume (as an indicator of tissue investment) when lower demands are placed on it [21–24]. We tested how neural investment related to both lifetime task repertoire differences between worker subcastes, and differences in modalities used by soldiers performing discrete defense tasks.

**Figure 1.**
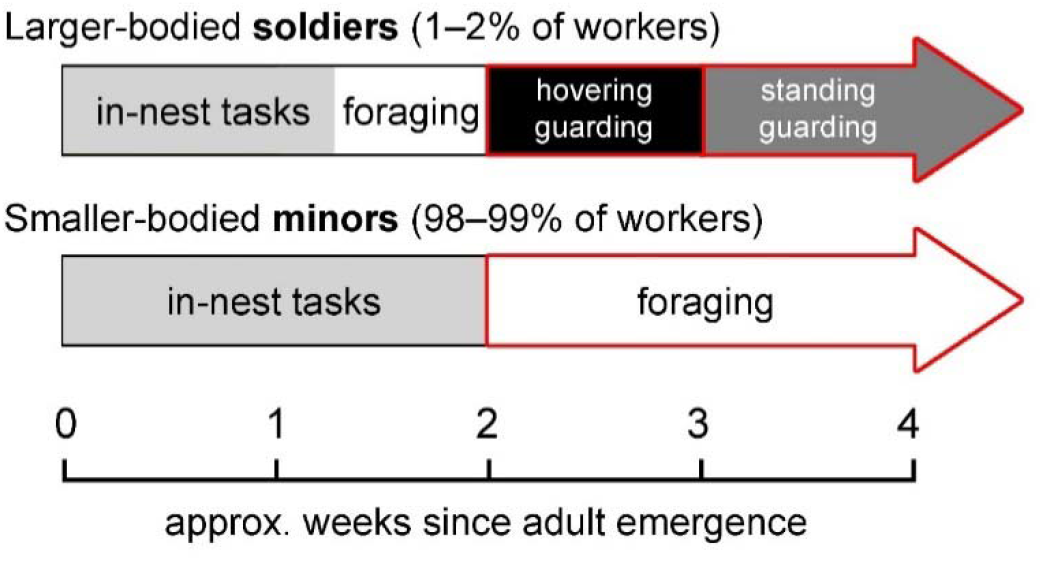
Adult age trajectories of soldiers versus minors within colonies of *Tetragonisca angustula*, based on Hammel et al. [13] and Baudier et al. [14]. Red boxes show sampled focal task groups in this study. Foraging minors were collected according to behavior, not morphotype. This is possible because soldiers typically make up a small percentage of worker forces in general [12, 51] and likely make up an even smaller portion of foragers due to the shorter time they are believed to perform this task relative to minors [14]. As such, the estimated < 3% of accidentally collected foraging soldiers among foraging minors was expected to be negligible, as was confirmed by size measurements (Figure 3B).

Soldier ants with smaller lifetime task repertoires than minors have previously been shown to have a lower ratio of total brain volume to body size and a smaller portion of their brains consisting of mushroom bodies (regions associated with higher cognition) [3, 25]. However, eusocial insect soldiers do not always have smaller lifetime task repertoires than other workers. *Tetragonisca angustula* soldiers work 34% to 41% more than non-soldier nestmates and have a lifetime task repertoire size that is 23% to 34% larger [13]. Soldiers of *T. angustula* also specialize and sub-specialize on defense tasks while minors do not typically perform colony defense [13] unless in a crisis [14]. By comparing brain volume and mushroom-body volume between soldiers and minors of *T. angustula*, we also tested whether the widely observed trends of smaller relative brain and mushroom-body size in eusocial insect soldiers is observed for stingless bee soldiers, despite soldiers in this species actually having a larger lifetime task repertoire. Such a result would challenge the notion that repertoire size differences drive the relatively smaller size of soldier brains in social insects at large.

We also investigate within-soldier neural diversity associated with discrete defense tasks performed by stingless bee soldiers of different ages. By comparing neural investment in brain regions associated with visual versus olfactory acuity and processing, we investigate the degree of sensory-specific neural specialization which underpins this rapid age progression through discrete defense tasks. Soldier temporal polyethism across distinct defense tasks has only been demonstrated in two eusocial species previously [14, 26] and is still largely unstudied. To our knowledge, this is the first study to investigate the potential rapid neural investment transitions that underlie age-related shifts in in discrete defense tasks.

In this study, we simultaneously explore neuroanatomical correlates of the two most prominently studied forms of division of labor in social insect colonies: morphological castes and temporal polyethism. Together these cross-age and cross-morphotype comparisons of stingless bee neural investment inform a broader framework for considering underlying neuroanatomical characteristics of finely tuned but flexibly specialized individuals within heterogeneous social groups.

## Methods

### Field samples

We collected all bees from naturally occurring nests in the town of Gamboa in Colón Province, Panama (9° 7’ N, 79° 42’ W). Per established methods [13, 14, 19], at each nest entrance, we collected bees from three discrete task groups (Figure 1): standing guards (typically 2- to 3-week-old soldiers), hovering guards (typically 3- to 4-week-old soldiers), and foragers (predominantly 2- to 4-week-old minors). We selected minors while in the task of foraging to compare to soldiers for total brain investment comparisons because they were of similar age but contrasting morphotype. A bee was deemed a forager if it exited the nest and immediately flew away from the nest entrance (distinguished from waste-removal workers by lack of carried detritus). We observed nest entrance guards for 20s each. If a bee spent the full duration of that time standing motionless on the nest entrance tube while facing toward the entrance of the tube, we considered it a standing guard. If a bee spent the full 20s flying in static formation in front of the nest entrance tube while facing inwards towards the flyway, we considered it a hovering guard.

We collected a total of 41 bees from 5 colonies live into 95% ethanol in February of 2018 for use in the assessment of neuroanatomical volumes (14 hovering guards, 13 standing guards, 14 foragers). In February 2019 we collected 50 additional bees (17 hovering guards, 16 standing guards, 17 foragers) from 5 colonies for immunohistochemistry. We anesthetized bees to be used for immunohistochemistry via 5-minute exposure to -40 °C before heads were separated from bodies. We then removed the caudal cuticle and muscle of the head before fixing and storing the brains in 4% paraformaldehyde (PFA) in phosphate-buffered saline (PBS) at 4 °C.

### Volumetric histology and quantification

Following field collection, we photographed the heads of each ethanol specimen using a DSLR mounted camera on a dissecting scope with a micrometer, and quantified head width and height using ImageJ 1.52 digital imaging analysis software [27]. We approximated head capsule volume from head width and height, assuming an ellipsoid shape with head depth equal to half of head height.

We stored heads in Prefer glyoxal fixative (Anatech Ltd) for a minimum of 7 days before dehydrating them in a series of increasing ethanol, acetone, then Embed 812 plastic resin concentrations (Electron Microscopy Sciences). We incubated heads in 0.1 ml resin at 60 °C for 72 hours in pyramid molds. We then cut resin embedded heads along the frontal plane into 10 um or 7 um thick sections using a rotary microtome, mounted them on gelatin-coated microscope slides, stained them with Toluidine blue (stain was cleared in a series of distilled water, increasing ethanol concentrations, and HistoChoice^®^ clearing medium), and cover-slipped them under DEPEX transparent mounting medium.

We photographed every section containing brain tissue using a compound light microscope-mounted digital camera at 5x magnification using LAS V4.9 software (Figure 2A). We quantified brain region area in ImageJ, and multiplied area by section thickness to estimate slice volume, summing the volumes across all slices to estimate total brain region volumes. We used Reconstruct Software (https://synapseweb.clm.utexas.edu/software-0, April 2020) to generate 3D reconstructions of the brain regions quantified: mushroom bodies (calyx lip, calyx collar, basal ring, peduncles), antennal lobes, optic lobes (medulla and lobula only), central complex, and the rest of the brain (protocerebrum, protocerebral bridge, subesophageal ganglion) (Figure 2B).

**Figure 2.**
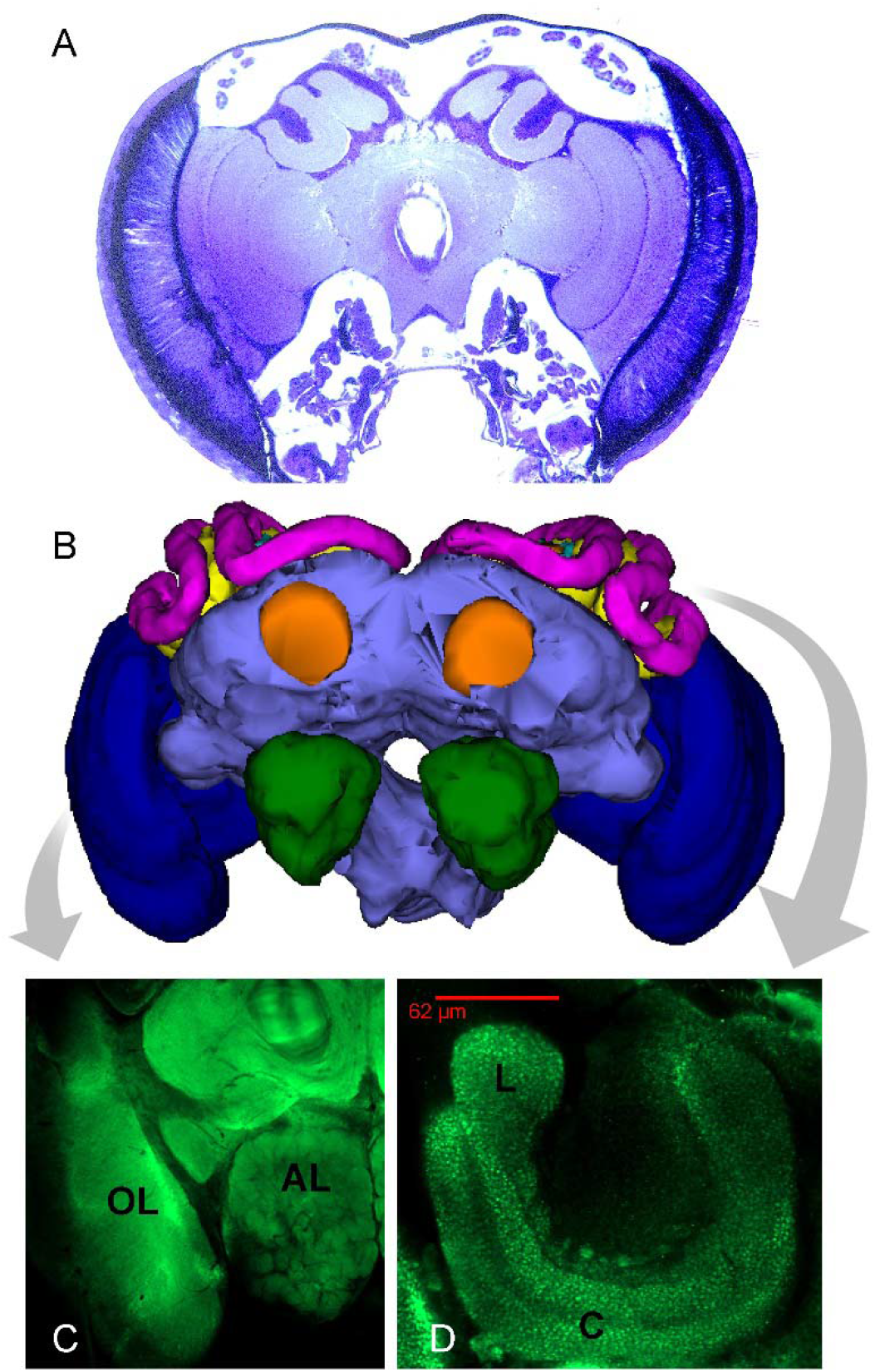
**(A)** A representative brain section from resin-embedded slicing histology done to quantify brain region volumes. **(B)** 3-dimensional reconstruction based on volumetric measurements of resin-embedded specimens. Dark blue = optic lobe, green = antennal lobe, magenta = lip, yellow = collar, aqua = basal ring, orange = peduncle, lavender = central brain mass. **(C)** Synapsin immunostaining of peripheral sensory regions of interest: optic lobe (OL) and antennal lobe (AL). **(D)** Synapsin immunostaining of sensory regions of interest within the mushroom body calyx: lip (L, olfaction) and collar (C, vision). Note visibly distinct synaptic vesicles.

All analyses in this study were conducted in R statistical software version 4.0.0. We tested for differences in relative volumes (region volume / total brain volume) of peripheral visual sub-regions (optic lobes) [28], peripheral olfactory sub-regions (antennal lobes) [29], central visual processing (collar) [30], and central olfactory processing (lip) [30] across task groups (minors collected while foraging, hovering guard soldiers, and standing guard soldiers) using separate mixed-effect ANOVAs that took into account colony as a random factor. See also the supplementary information (Figure S1) for comparison of all measured region volumes at large using a Principal Component Analysis.

Total brain volume as well as relative volume of mushroom bodies (brain regions associated with higher cognition) are commonly used metrics for neural tissue investment in social insects [1, 25, 31, 32]. To test whether investment is better predicted by bee size or lifetime task repertoire size, we compared absolute and relative total brain volume and mushroom-body volume across soldiers (pooled hovering guards and standing guards) and minors. First, to confirm that collected morphotypes (minors versus soldiers) were different in size as expected, we fit a linear mixed-effect model which included colony as a random factor, subcaste (soldier vs minor) as a fixed factor, and head width as a response variable. We compared absolute brain sizes by fitting a linear mixed-effect model that included colony as a random factor, subcaste as a fixed factor, and absolute total brain volume as a response variable. We compared relative brain volumes by fitting a liner mixed-effect model that included colony as a random factor, subcaste as a fixed factor, and relative brain volume (brain volume / head capsule volume) as a response variable. Separate Type II Wald Chi-square tests were used to test for predictive significance of morphological subcaste (soldiers versus minors) in all three models. We used a mixed-effect ANCOVA (colony was a random factor) to test for differences between morphological subcastes in the relationship between mushroom-body volume and total brain volume. (log-transformed data are also presented in supplementary material, Figure S2).

### Immunohistochemistry and synaptic quantification

Partially dissected brains were fixed in 4% PFA at 4 °C for 7 months, after which they were completely removed from the head capsule and subjected to synaptic staining histology. We used a modification of an existing immunohistochemistry protocol [33, 34] with increased incubation times to allow for improved penetration in these stored specimens. Brains were made permeable by washing (4×20 min) with 0.5% Trition-X100 phosphate-buffered saline (PBS-T) while on a shaker. After PBS-T was removed, the samples were blocked with 4% Normal Goat Serum (NGS) and placed on a shaker for 1 hour at room temperature. Following the removal of NGS, the primary antibody SYNORF1 was added in a 1:10 dilution of 0.5% PBS-T for 7 days at 4 □. After another PBS-T wash (4×15 min) the brains were incubated with the secondary antibody, 488 anti-mouse 1:50 in 4% NGS, for 7 days at 4□. Brains were mounted with antennal lobes oriented upward using Vectashield^®^ mounting medium (Maravai Life Sciences, San Diego, CA).

We imaged whole mount preparations using a Zeiss LSM800 confocal microscope (Carl Zeiss Vision Inc., San Diego, CA), using an argon laser at 448 nm. Optical sections had a resolution of 1024 x 1024 pixels. Five representative section images were taken of each brain: a whole-brain image at 10X taken at a depth approximate to the centroid of the antennal lobes (Figure 2C) for quantifying antennal and optic lobes, and four 20X images, one of each mushroom body calyx, taken at the shallowest depth at which the lip separated and the collar was visible in frame (Figure 2D) for quantifying lip and collar synaptic staining. Gain and digital contrast were adjusted in each image to ensure that none of the focal regions of interest (antennal lobes and optic lobes for whole brain images, lips and collars for mushroom-body images) were over or underexposed.

We assessed brightness in each slice image using ImageJ by tracing the edges of the lip, collar, antennal lobe and optic lobe (medulla and lobula), then calculating the mean pixel intensity (brightness) of each region. For antennal and optic lobes, we quantified both right and left sides, but in a minority of cases a side of the brain that was damaged during dissection was omitted from the study (N = 86 imaged). In the majority of cases, all 4 mushroom body calyces (and 4 lip/collar synaptic density estimates) were imaged and quantified. However, calyces that were damaged during dissection or did not stain sufficiently to make imaging possible were omitted from analysis (N = 146 imaged). Brightness ratios were calculated for olfactory versus visual regions within each slice image (lip versus collar or antennal lobe versus optic lobe, averaged across medulla and lobula). Brightness ratios were log transformed to normalize distributions. We compared the log_10_ of the ratio of lip to collar brightness across mushroom bodies within hovering guards, standing guards, and foragers using a mixed-effect ANOVA that took into account Colony ID and Bee ID as nested random factors. We compared the log_10_ of the ratio of antennal lobe to optic lobe brightness across sides of the brain in hovering guards, standing guards, and foragers using a mixed-effect ANOVA that took into account Colony ID and Bee ID as random factors.

We estimated synaptic density of lip (olfactory) and collar (visual) calyces from representative slice images of each mushroom body calyx, using the Image-based Tool for Counting Nuclei (ITCN) plugin in ImageJ [35]. Images were inverted and converted to 8-bit before ITCN estimated number of peaks per μm^2^. Based on preliminary image measurements, target peak width was 10 pixels and minimum peak distance was 5 pixels. This was done only for mushroom body calyx images. The ratio of detected synaptic vesicles per μm^2^ (lip/collar) was compared across hovering guards, standing guards, and foragers using a mixed-effect ANOVA that took into account Colony ID and Bee ID as nested random factors. Absolute detected synaptic vesicles per μm^2^ was compared across hovering guards, standing guards, and foragers using two separate mixed-effect ANOVAs (one for lip, one for collar) that took into account Colony ID and Bee ID as nested random factors. In the majority of cases, 4 mushroom body calyces (and 4 lip/collar synaptic density estimates) were imaged and quantified. However, calyces damaged during dissection or that did not stain sufficiently to make imaging criteria possible were omitted from analysis (N =166 imaged).

## Results

### Brain volumes

Bees with larger lifetime task repertoires (soldiers) did not have greater total brain volume or investment in higher-cognition regions (mushroom bodies) compared to smaller lifetime task repertoires (minors). Absolute brain volume did not differ between soldiers and minors (*X^2^* = 1.44, *p* = 0.231, Figure 3A) despite soldiers having significantly larger heads (*X^2^* = 17.27, *p* < 0.001, Figure 3B). The portion of head capsule occupied by brain tissue was therefore higher in minors than in soldiers (*X^2^* = 5.22, *p* = 0.022, Figure 3C). Although higher cognition regions of the brain (mushroom bodies) increased linearly with total brain size, soldiers and minors did not differ in mushroom-body volume or in the slope of the relationship between mushroom-body volume versus total brain volume (ANCOVA; Table 1, Figure 3D). Head volume was also generally a poor predictor of absolute brain volume (linear regression, *R*^2^ = 0.07, F_1,39_ = 3.99, *p* = 0.053; Figure 3E).

**Figure 3.**
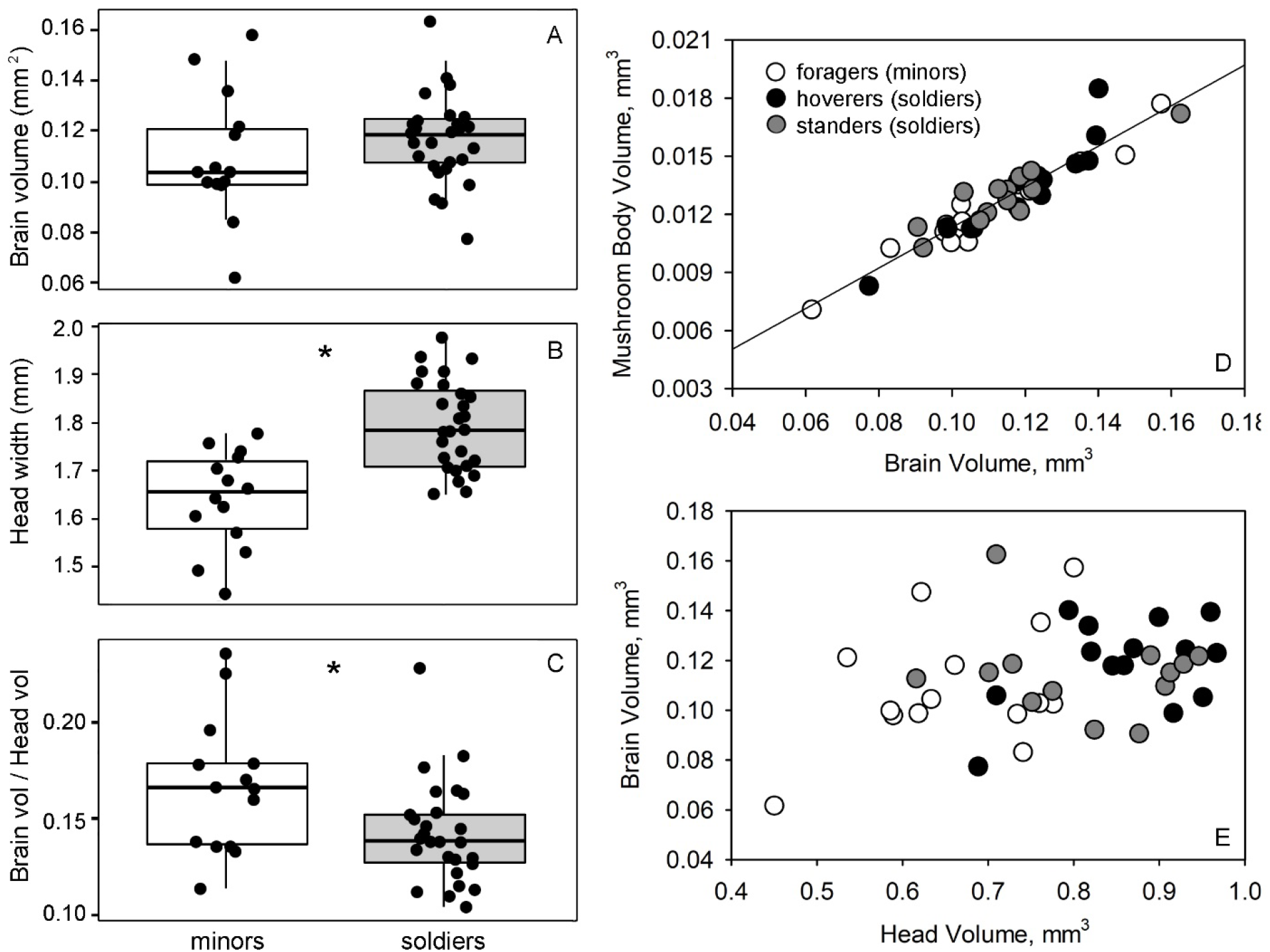
External and internal morphometric differences between distinct worker subcastes (minors versus soldiers) of specimens sampled for volumetric data collection (N = 41). * indicates significant difference between soldiers and minors (α = 0.05). **(A)** Total brain volume of soldiers vs. minors. **(B)** Difference in head width between soldiers and minors. **(C)** Difference in portion of the head capsule composed of brain tissue between soldiers and minors. **(D)** Mushroom body (lip + collar) volume increasing linearly with total brain volume, regardless of morphotype or specific soldier task group (Table 1). **(E)** Lack of a relationship between brain volume and head capsule volume, with point color representing morphotype and task group.

**Table 1.** Statistical output of mixed-effect ANCOVA, with a model of structure: Mushroom_body_volume ~ Brain_volume * Morphotype + (1|Colony), where morphotype is a categorical variable with two levels (soldier and minor).

| Fixed factors | $X^2$ | $df$ | $P$ |
| --- | --- | --- | --- |
| Brain volume | 385.91 | 1 | < 0.001 |
| Morphotype | 0.58 | 1 | 0.446 |
| Brain volume : Morphotype | 1.11 | 1 | 0.291 |

Relative optic lobe volume significantly differed across task groups (*X^2^* = 5.99, *df* = 2, *p* = 0.049; Figure 4A). Compared to minors, soldiers had a larger portion of their brains dedicated to optic lobes but only while in the visually demanding task of hovering guarding (*z* = 2.42, *p* = 0.041). Bees that had already aged into the task of standing guarding had similar relative optic lobe volumes as minors (*z* = 0.86, *p* = 0.667) but also did not significantly differ from hovering guards (*z* = -1.545, *p* = 0.270). Standing guards, hovering guards, and foraging minors had the same percentage of brain volume dedicated to antennal lobe (*X^2^* = 4.83, *df* = 2, *p* = 0.089; Figure 4B), mushroom-body collar (*X^2^* = 0.844, *df* = 2, *p =* 0.656; Figure 4C), and mushroom-body lip (*X^2^* = 0.69, *df* = 2, *p* = 0.708; Figure 4D). Proportions of regional brain volumes were otherwise quite similar across task groups, as indicated by Principal Component Analysis (Supplementary Figure S1).

**Figure 4.**
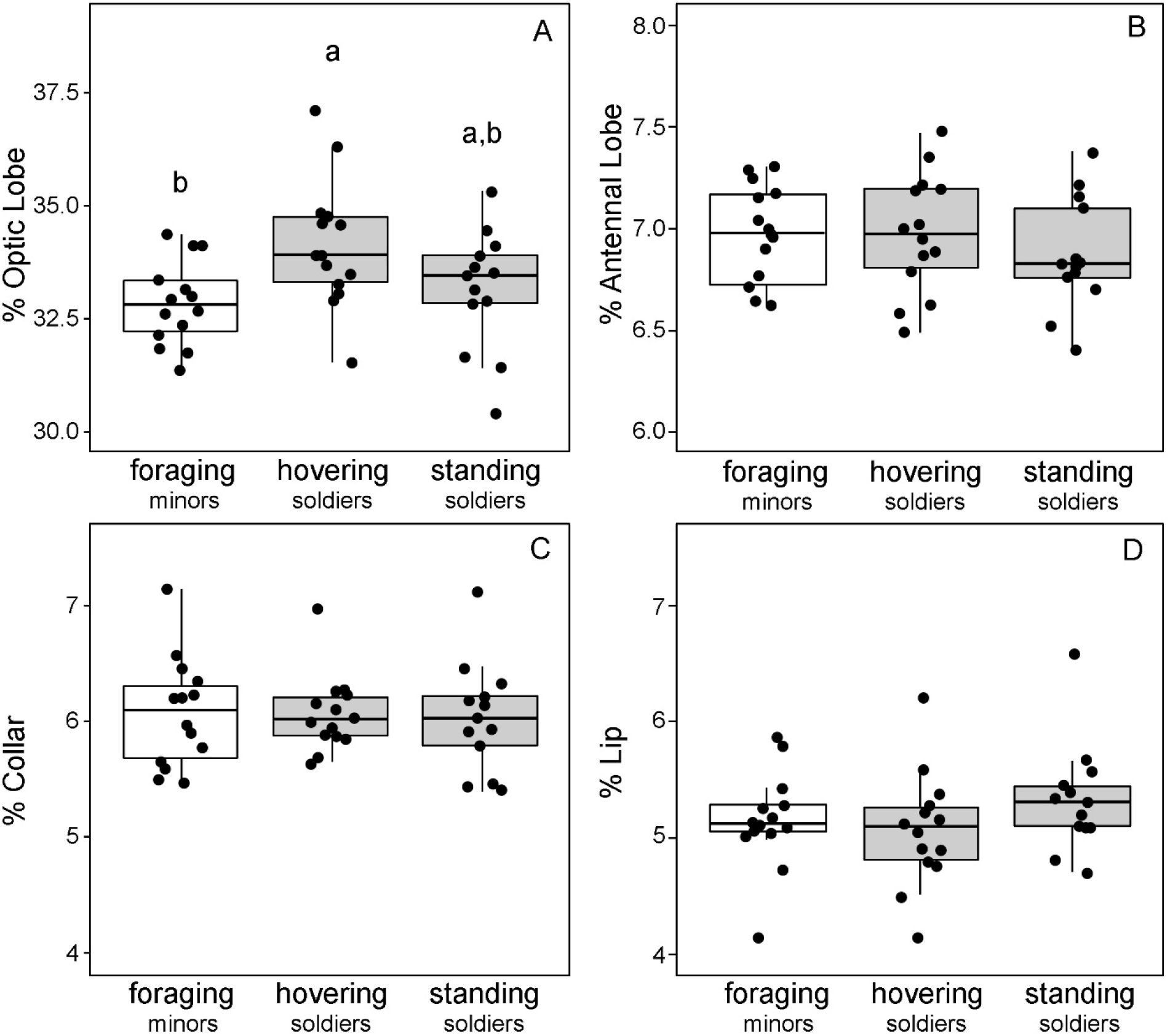
Comparisons of relative investment (percent of brain volume) comprised by peripheral brain regions associated with vision (optic lobe, **A**) and olfaction (antennal lobe, **B**), as well as sensory processing regions associated with vision (Collar, **C**) and olfaction (Lip, **D**). In all cases N = 41. Grey boxes denote defense tasks performed by morphologically larger soldier bees. White boxes denote a task (foraging) performed predominantly by small-bodied minors (see Figure 3B). Lowercase letters denote results of Tukey HSD post hoc tests performed for regions that differed in relative volume according to linear mixed-effect analyses (colony = random factor).

### Synaptic brightness and density

Small-bodied foragers and large-bodied hovering and standing guards did not significantly differ in the proportionate synaptic density of olfactory regions (lips) versus visual regions (collars) of the mushroom bodies (*X^2^* = 0.23, *df* = 2, *p* = 0.889; Figure 5A). These three groups also had the same ratio of synaptic brightness between lip and collar (*X^2^* = 1.52, *df* = 2, *p* = 0.468; Figure 5B) and between peripheral olfactory regions (antennal lobes) and peripheral visual regions (optic lobes) (*X^2^* = 1.92, df = 2, *p* = 0.382; Figure 5C). There were also no significant differences among these three groups in absolute synaptic density within the lip (*X^2^* = 4.56, *df* = 2, *p* = 0.102) or collar (*X^2^* = 5.33, *df* = 2, *p* = 0.070).

**Figure 5.**
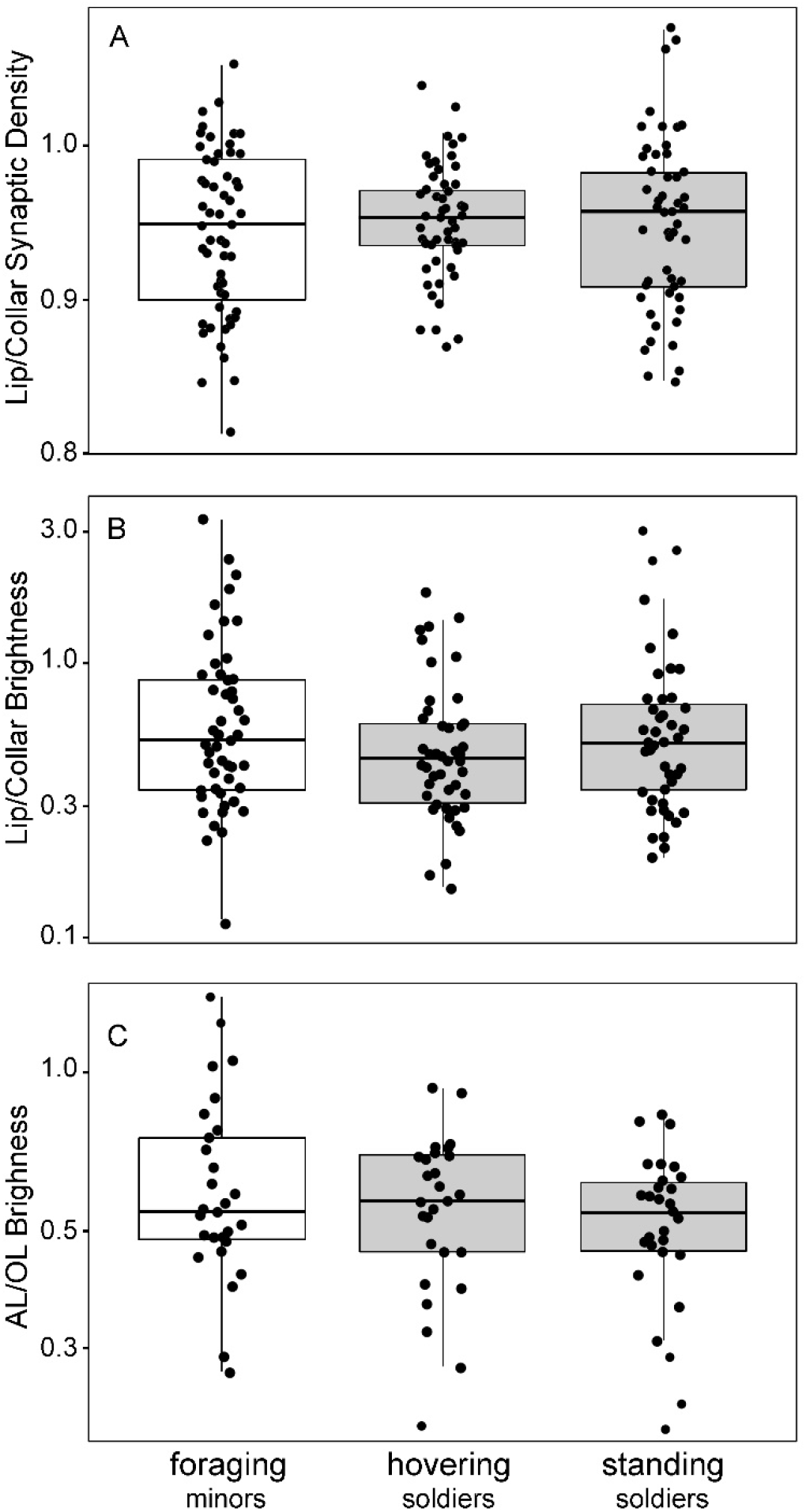
Quantified synapsin in brain regions associated with olfactory versus visual sensing and processing. Grey are defensive task groups performed by morphologically larger-bodied soldiers. Meanwhile foraging is performed predominantly by smaller-bodied minors of similar age **(A)** Ratio of lip (olfactory processing) to collar (visual processing) estimated synaptic density did not differ across groups; N =166. **(B)** Log-transformed ratio of lip to collar brightness (pixel density) did not differ across groups; N =146. **(C)** Log-transformed ratio of antennal lobe (AL) brightness to optic lobe (OL) brightness did not differ across groups; N = 82.

## Discussion

### Total brain investment

Soldier bees with larger lifetime task repertoires had similar absolute brain volumes, and smaller ratios of brain volume to head volume, compared to similarly aged minors with smaller lifetime task repertoires. Soldiers and minors did not have different relative mushroom-body volumes, regions associated with high cognition [1, 34, 36]. Overall, these results do not support the hypothesis that contrasts in lifetime task repertoire drive differences in brain size between soldiers and non-soldiers. However, we did observe that stingless bee soldiers had smaller relative brain size to head size, which is similar to patterns observed in army ants [25] and turtle ants [3]. Together, this suggests that factors other than lifetime task repertoire size likely explain contrasts in relative brain size between soldiers and other workers within social insect colonies.

These results raise the important question of which timescales of task repertoire size are most relevant to total neural investment. Previous studies of soldier brain size used species for which both instantaneous and lifetime task repertoire size are generally smaller in soldiers than in other workers. However, task repertoire size is complex and dynamic for *T. angustula* soldiers. *Tetragonisca angustula* soldiers work 34% to 41% more than non-soldier nestmates and have a lifetime task repertoire size that is 23% to 34% larger, but much of this task diversity occurs early in a soldier’s adult life [13]. Whether and to what extent the shift to primarily defense tasks towards the end of a soldier’s life coincides with a reduction in neural investment remains an interesting and relevant question. Such a reduction in total brain size coinciding with a decrease in task repertoire size has been reported in harvester ant queens following nest founding [37] and in *Harpegnathos saltator* as they reversibly shift from foraging to reproductive (gamergate) tasks [38].

Another striking possibility is that coinciding increases in muscular investment within soldier bodies to enable improved combat capabilities affects relative brain size, a notion originally proposed by O’Donnell et al. [25]. Similar to the ant soldiers for which this was originally posited, stingless bee soldiers also defend the colony from intruders using biting [12, 39], which places a demand for higher muscular investment within the head capsules of soldiers. How muscular and neural investment needs interplay among individuals within a coordinated social group is still largely unexplored.

Lastly, it is possible that the broadly observed trend for relatively larger brains among smaller colony members in eusocial Hymenoptera is due to the physiological scaling constraints of neural tissue more than either task repertoire or muscular investment. Haller’s rule predicts larger relative brain size in smaller-bodied animals via negative allometry between body and brain volumes [40, 41]. Consistent with Haller’s rule we did find that smaller bees within the colony had relatively larger brains, however the relationship between head size and brain size was not significant, and so we did not find evidence of negative allometry.

### Modality-specific volume contrasts

Soldiers had different modality-specific neural investment than minors but only at certain ages and according to task demands on that age group. Compared to smaller-bodied nestmates of similar age, soldiers had significantly greater relative optic lobe size, but only while engaged in the more visually demanding defensive task of hovering guarding. As soldiers aged into the less visually demanding task of standing guarding, this difference in peripheral visual investment between soldiers and minors disappeared. Notably, these modality-specific differences seemed to be primarily in peripheral brain regions associated with visual acuity (optic lobe) and not visual information processing (mushroom body collar) [30]. Similar peripheral brain investment contrasts have been observed across dimorphic male bees employing alternative reproductive tactics in *Centris pallida* and *Amegilla dawsoni* [42] and between males and females of the sexually dimorphic longhorn bee *Eucera berlandi* [43]. However, here we report for the first time, similar patterns across age-differentiated soldier sub-castes. The transition from hovering guarding to standing guarding in *T. angustula* soldiers occurs over approximately seven days [14]. Although the magnitude of observed optic lobe volumetric change is small, and only significant relative to optic lobe volume of minors, the rapid pace of this transition is striking and suggests the importance of future investigations of the neurobiology of soldier sub-specialization in other eusocial taxa.

We also report no differences (peripheral or otherwise) in volumetric olfactory investment across minors versus soldiers or hovering versus standing soldiers. Although standing and hovering guards differ in which odors elicit responses [19], both hovering and standing guarding are tasks that demand some degree of olfactory intruder detection [16, 44]. Our findings of similar gross olfactory investment between hovering and standing guards is consistent with these possibly similar, but difficult to compare, demands on olfactory acuity and processing.

Because hovering guards and standing guards differ not only in tasks performed but also in age (ca. 7 days), whether and how experience versus experience-independent ontogeny mediate the subtle neural transition (decrease) in optic lobe remains an open question. Contrary to the patterns we report, optic investment increases with both age and visual experience in young adult *Drosophila melanogaster* [45] and *Polistes fuscatus* [46]. However, soldiers typically perform standing guarding tasks in their final week of life (Figure 1) [13], and so a relationship between volumetric change and senescence may be more likely. Optic lobes of the Europrean honey bee (*Apis melifera*) have been found to be especially susceptible to oxidative stress associated with senescence [47]. Whether similar processes relate to the patterns in brain volumetrics we report here remains an open question.

### Synaptic density and brightness

Synaptic activity is remarkably plastic and can change in social hymenopteran brains throughout adulthood due to age or life experiences, independent of shifts in neuropil volume [48]. Indeed, despite observing contrasts among task groups in total and modality-specific brain volume, we found no evidence of differences in the ratio of modality-specific synaptic density or brightness across task groups. In honey bees, increases in synaptic (microglomerular) density within mushroom bodies have been associated with the formation of long term modality-specific memory [49]. Among *T. angustula*, standing guards are, in a sense, older and more experienced hovering guards with potentially greater accumulated long-term memories. However, we report no evidence of synaptic density differences between standing and hovering guards in either lip or collar using these methods. These results are contrary to our prediction and may indicate that brain volume but not synaptic activity drive differences in these focal behaviors. Other mechanisms involved in energy metabolism that were not measured in this study could also help explain differences in behavioral phenotypes. Beyond ATP production, mitochondria play a role in maintaining intracellular calcium homeostasis, which is critical for cell signaling [50]. Whether and how task groups differ in neural mitochondrial density and function within this study remains an open question.

### Conclusion

In exploring the neurobiology of stingless bee soldiers, we report both contrasts and similarity in neural architecture across highly sub-specialized but behaviorally flexible stingless bee workers. Although both age and morphological worker sub-caste were predictive of some aspects of regional and total brain investment, we also observed striking overlap in brain region volumes at large (Supplementary Figure S1) and similarity in synaptic/microglomelular density. The neural architecture of worker castes of *T. angustula* is quite similar, though mildly specialized across discrete task-groups, and is consistent with this species’ documented investment in simultaneous strategies for specialization but also flexibility in defensive task allocation [14]. Although the vast majority of guarding tasks are performed by soldiers under normal circumstances, minors are capable of performing some degree of nest defense when soldiers are removed [14]. Neural architecture that is similar but slightly specialized likely enables both defense specialization, and also flexible defensive reallocation in a crisis. As the first study of neural architecture of morphologically distinct soldier brains in a bee, these results broaden our understanding of the neurological mechanisms underlying defense specialization in highly coordinated societies.

## Acknowledgements

We thank the Smithsonian Tropical Research Institute and the community of Gamboa, Panama for allowing access to public and private lands on which this field work was conducted. Permits for this work were issued by the Panamanian Ministry of the Environment (MIAMBIENTE). Funding was provided by the Arizona State University, School of Life Sciences, Innovative Postdoctoral Research Award to MMB and KMB, and by contract W31P4Q-18-C-0054 from the United States Defense Advanced Research Projects Agency (DARPA). We thank Hermógenes Fernández-Marin, David Roubik, and Madeleine Ostwald for field assistance, and thank Purnima Sachdeva and Rheanna Congdon for lab assistance. We thank Majid Ghaninia Tabarestani, Jason Newbern, Brian Smith, and Jon Harrison for helpful discussion of methodology and use of facilities and equipment.

## Notes

### Competing Interest Statement

The authors have declared no competing interest.

