## Supplemental information for "Soldier neural architecture is temporarily modality-specialized but poorly predicted by repertoire size in the stingless bee *Tetragonisca angustula*"

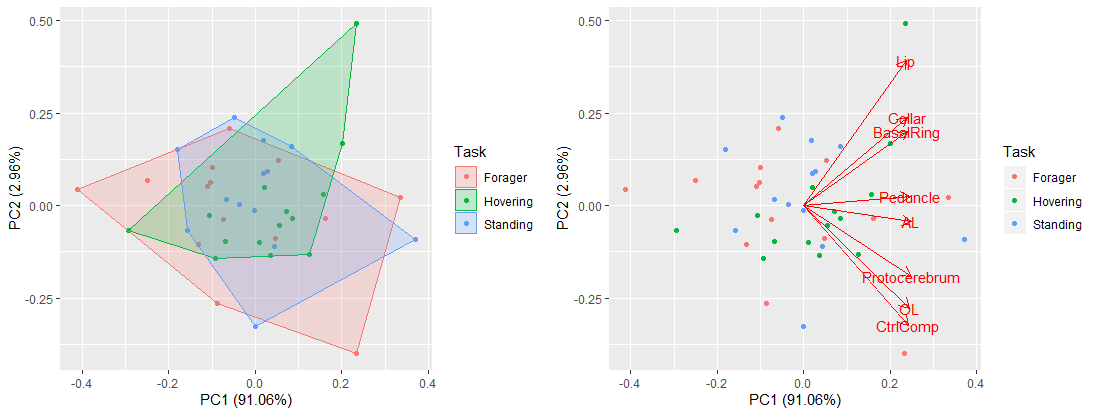

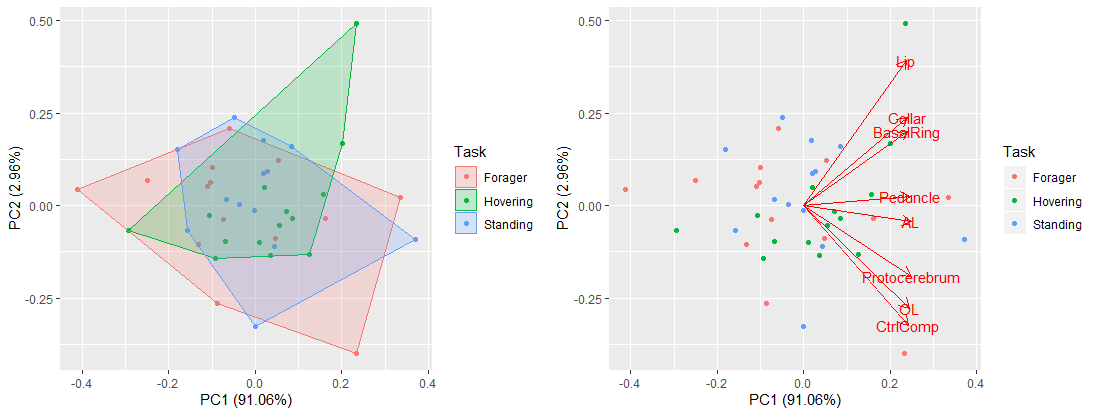

**Supplementary Figure S1**. Principal component analysis of all measured brain regions, with color showing contrasts of focal task groups in this study. AL = Antennal Lobe, OL = Optic Lobe, CrtlComp = Central Complex.

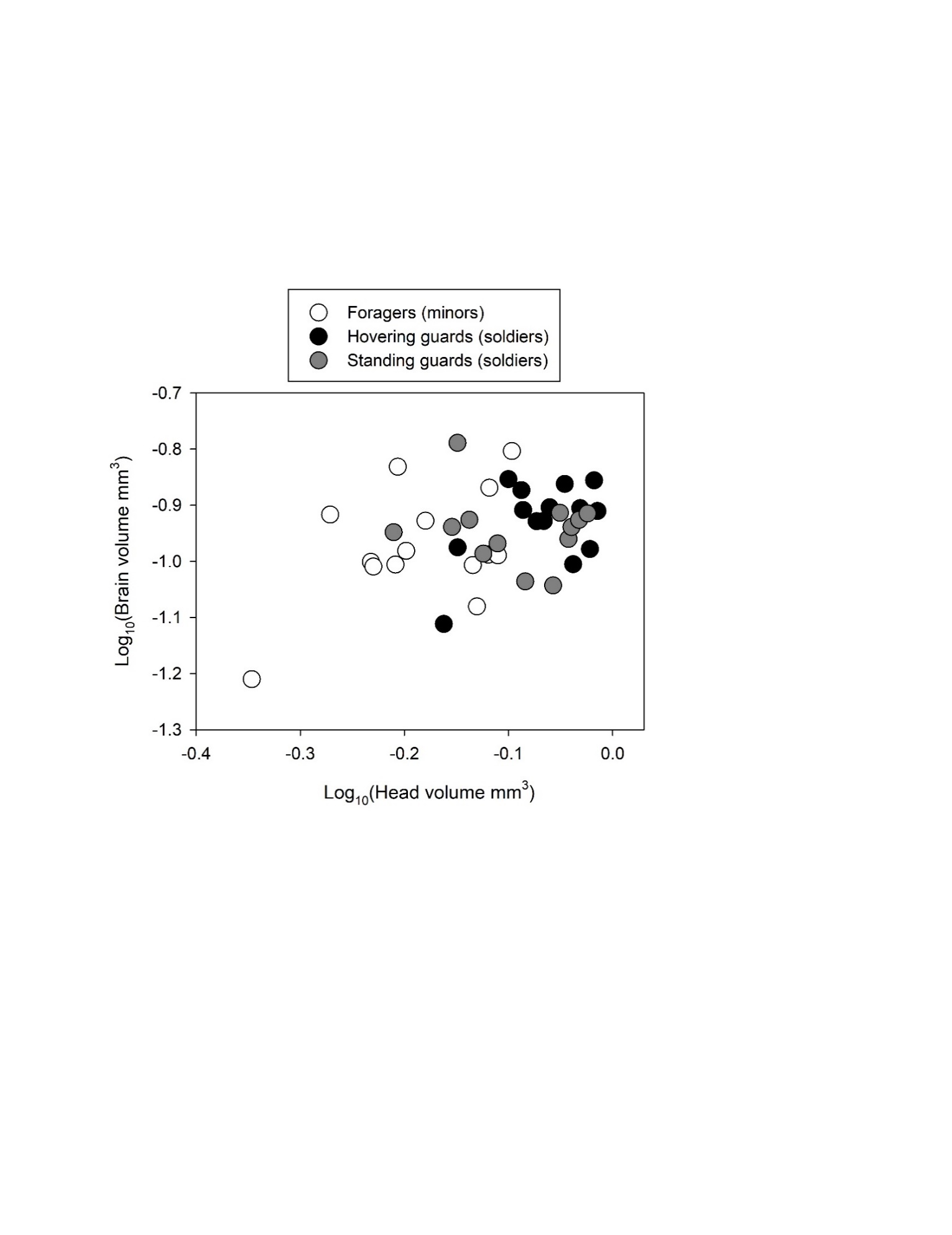

**Supplementary Figure S2.** Log-log transformed head volume versus brain volume with point colors denoting morphological and task groups. Log-transformed data were not used in the main analyses because such transformations increased heteroskedasticity. However, transformed and untransformed data suggest the same lack of predictability of total worker bee brain size based on head volume.

**Supplementary datasets to be submitted upon article acceptance**

**Supplementary Data S1.** File name “Data_S1_volumes.csv”. All brain volumetric data and accompanying morphological data analyzed in this study.

**Supplementary Data S2.** File name “Data_S2_synaptic_brightness.csv” All synaptic brightness (intensity) data analyzed in this study.

**Supplementary Data S3.** File name “Data_S3_synaptic_density.csv” All synaptic density data analyzed in this study.
